# Changes of mind after movement onset: a motor-state dependent decision-making process

**DOI:** 10.1101/2021.02.15.431196

**Authors:** Ignasi Cos, Giovanni Pezzulo, Paul Cisek

## Abstract

Decision-making is traditionally described as a cognitive process of deliberation followed by commitment to an action choice, preceding the planning and execution of the chosen action. However, this is challenged by recent data suggesting that multiple options are specified simultaneously and compete in pre-motor cortical areas for selection and execution. Previous studies focused on the competition during planning, and leave unaddressed the dynamics of decisions during movement. Does deliberation extend into the execution phase? Are non-selected options still considered? Here we studied a decision-making task in which human participants were instructed to select a reaching path trajectory from an origin to a rectangular target, where reward was distributed non-uniformly at the target. Critically, we applied mechanical perturbations to the arm during movement to study under which conditions such perturbations produce changes of mind. Our results show that participants initially selected the direction of movement towards the highest reward region, and changed their mind most frequently when the two choices offered the same reward, showing that deliberation continues and follows cost-benefit considerations during movement. Furthermore, changes of mind were dependent upon the intensity of the perturbation and the current state of the motor system, including velocity and distance to targets. Although reward remains most relevant, our results indicate that the state of the motor system when the perturbation occurs is a crucial determinant of changes of mind. This indicates that the neural circuits that assess reward and those that control movements operate synergistically rather than sequentially during decision-making.

**Significance Statement:** Our study provides supporting evidence for the notion that deliberation during decision-making continues after movement onset because unselected potential actions are not completely suppressed or discarded. From a neurophysiological perspective, our findings suggest that the competition between actions is not over before action initiation, possibly because the initially unselected neuronal population retains some sub-threshold activation, which enables them to take control afterwards. Furthermore, our findings also suggest that decision-makers have a variable degree of commitment to their initial choice, which depends on the relative reward of the offers and on the state of the motor system. The commitment is stronger if the initially selected plan leads to higher rewards, and changes of mind occur more frequently if the velocity and relative position of the end-point are within specific ranges.

## INTRODUCTION

Decision-making has been traditionally described as a cognitive process, which is completed prior to the preparation and execution of the action that reports the choice (Newell and Simon, 1972). However, this serial model was developed for *laboratory decisions*, where options are fixed and actions can be executed almost instantaneously. By contrast, the brain’s decision systems plausibly evolved to deal with *situated decisions* – for example, a lioness deciding which gazelle to chase – that pose different adaptive challenges (Filimon et al., 2013; Cisek and Pastor-Bernier, 2014a; Ratcliffe and Newport, 2017; Wispinski et al., 2020).

Waiting to reach a decision is rarely possible in situated decisions. Most often a movement must be started before completely making up your mind, not to miss important opportunities. This may be better fulfilled by brain architectures where decision and motor systems are more intertwined than traditionally assumed, with motor regions engaged early on during deliberation and possibly participating of the decision itself. Experimental evidence revealed that premotor cortex and superior colliculus may encode several competing potential actions prior to movement, and that the decision process can be characterized as a competition (McPeek and Keller, 2002; Cisek and Kalaska, 2010). This suggests that decisions may be performed by neural areas including sensorimotor regions, rather than being confined to prefrontal areas (Cisek, 2012; Kubanek and Snyder, 2015).

Involving motor cortices in decisions addresses another challenge of situated decisions: their continuous nature, since actions take time to complete. Furthermore, the environment is non-stationary and key determinants such as the geometric arrangement of targets (e.g., the distance to the prey) and action costs --- e.g., energy expenditure to reach a moving target (Morel et al., 2017), can change continuously as the action unfolds. Furthermore, new opportunities may become available or unavailable --- e.g., a novel prey can appear (Diamond et al., 2017; Michalski et al., 2020). Accordingly, animals have to continuously re-evaluate their choices after movement onset, rather than committing rigidly to their initial decisions (Lepora and Pezzulo, 2015). Recent studies have shown that movements may be initiated before the decision is complete (Wolpert and Landy, 2012), to later be revised, sometimes causing “changes of mind” (Resulaj et al., 2009; Song and Nakayama, 2009; Barca and Pezzulo, 2012).

In sum, this body of evidence suggests that a situated decision may be neurally implemented as a continuous competition between potential actions, and that spontaneous adjustments and “changes of mind” may occur after movement onset.

Interestingly, it has been shown that changes of mind are not just spontaneous but can also be triggered externally, by sudden target jumps, perturbations to the motor apparatus or changes of the environment (Nashed et al., 2012, 2014; Burk et al., 2014; Atiya et al., 2020; Marti-Marca et al., 2020). However, it is unclear whether these “externally triggered” changes of mind reflect truly deliberative processes based on cost-benefit considerations (Shadmehr et al., 2010; Rigoux and Guigon, 2012; Taniai and Nishii, 2015) or are simpler motor reflexes. Furthermore, if these changes of mind reflect deliberation, what information is considered? If they are influenced by economic variables such as the reward of the non-selected offer, then they should occur less often when the reward of the non-selected offer is lower than the selected option. If they take into account the momentary state of the motor system, then they should occur less often if the perturbation happens when the state of the motor system favours the selected offer, i.e., when one is close to the selected target and/or moving quickly toward it.

To investigate these questions, we designed a reward-driven reach decision task in which movements were sometimes perturbed, and predicted that changes of mind should occur more often with strong and early perturbations, when actions are slower, and when the arm position is further away from the initially selected target.

## MATERIALS AND METHODS

### Participants

A total of 16 subjects (6 males and 10 females, aged 21–36 years, all right hand dominant) participated in the experimental task. All subjects were neurologically healthy, had normal or corrected to normal vision, were naïve as to the purpose of the study and gave informed consent before participating. The study was approved by the Human Research Ethics Committee of the Faculté de Médecine, Université de Montréal (CERES-17-050), and conducted in accordance with the committee’s ethical standards.

### Task Apparatus

During the experimental session, the participants were seated facing the projection system with the right arm supported in a horizontal plane by the KINARM robotic exoskeleton (BKIN Technologies; Kingston, ON). The KINARM permits elbow and shoulder movements on the horizontal plane, as well as controlled mechanical perturbations to the upper and lower arm sections (Scott, 1999). The display of cues and hand-position feedback were presented to the subject by projection onto a mirror, and the arm and hand were occluded and never visible during the experiment. Custom written software controlled the stimulus presentation and task data collection of shoulder and elbow kinematic and kinetic variables at 1000Hz. Data from each session was transferred to a MySQL Community Server database (Oracle, Santa Clara, CA) for further analysis with custom-designed Matlab scripts (Mathworks, Natick, MA).

### Behavioural Task

To determine if changes of mind take movement-related factors into account, we designed a reward-driven decision-making task in which human participants had to perform a planar movement from an origin cue to a wide rectangular target, in which the reward for each trial depended on where on that target the endpoint landed. The distribution of rewards was bimodal, with peaks at the extremes and zero in the centre.

We introduce two hypotheses regarding changes of mind: First, that changes of mind are sensitive to the relative value of the offers. To test this, we compared three bimodal distributions, one with identical rewards (3 vs. 3) and two with different rewards (1 vs. 5 and 5 vs. 1) at the two extremes. We predicted changes of mind to occur more often in the former case, in which there is no advantage for reaching to either side. Second, we predicted that changes of mind are influenced by the momentary state of the motor system (i.e., velocity and distance from alternative targets when the perturbation occurs); to this end, we applied mechanical perturbations perpendicular to the direction of movement (left or right), in approximately ¼ of the trials. These were applied at two different levels of intensity (weak or strong perturbations) and at two different time points during the movement (early or late perturbations). We predicted changes of mind to occur more often with strong and early perturbations, when actions were slower, and when the arm position was farther away from the initially selected target.

The task consisted of 720 trials, performed in a single session. In each trial, the participant was asked to perform a reaching movement from a circular origin cue (diameter: 1cm) to a wide rectangular target (width: 10cm; depth: 1cm), placed 15 cm away and rotated 135 degrees counter-clockwise (FIG 1A). The target’s location and rotation were chosen as to equalize the potential influence of motor costs for movements towards the right/left part of the rectangle, as it coincides with the direction of the arm maximal inertia (Hogan, 1985; Cos et al., 2011). In brief, movements towards either side of the target incurred approximately the same biomechanical cost. The goal for the subject was to maximize reward by aiming at specific positions along the long side of the rectangle. Since our goal was to assess the influence of reward expectations, at the beginning of each trial, the reward distribution was indicated to the subject using triangle displays for one of three bimodal distributions: 3-3, 1-5, 5-1 (FIG 1A). These distributions peaked at the right/left edges of the rectangle’s long edge, and decreased towards zero when approaching its center. The distribution was zero off the right/left sides of the rectangle, implying that reaching movements missing the target would be awarded zero reward. The subject’s instruction was to freely select a reaching movement from the origin to any position along the long side of the rectangle, and the reward obtained on each trial was contingent upon arrival position and the distribution of reward.

**Figure 1.**
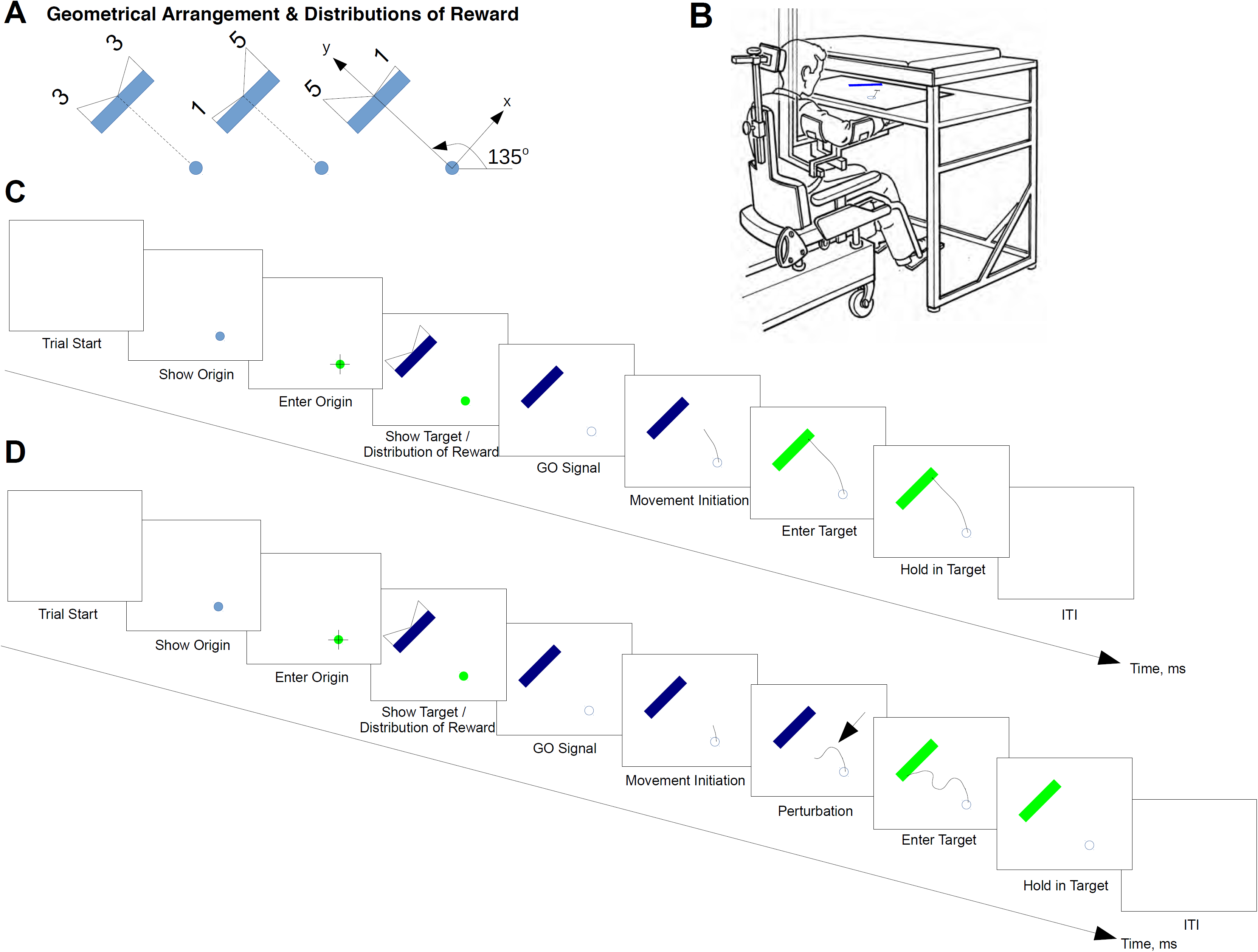
**A.** Geometrical arrangements of the stimuli on different trials consist of a circular origin cue (1cm) and a rectangular target (10cm wide, 1cm deep), placed about 15cm away from the origin at an orientation of 135 degrees. We show the three distributions of reward value, from left to right 3-3; 1-5; 5-1. respectively. **B.** KINARM setup. **C.** Time-course of a baseline trial. The trial starts with an empty screen for 1000ms. After this interval, the origin cue (1cm blue circular cue) is shown on the bottom-right part of the screen. When the endpoint (right-hand fingertip) enters the cue, the cue turns green, and the rectangular target and the distribution of value are shown on the top-left part of the screen. One second later the origin cue turns white to indicate the GO signal. If the subject’s endpoint leaves before the GO is given, the screen turns blank and the trial is invalidated. When the endpoint enters the target, the rectangular cue turns green. The screen turns blank 500ms after that. **D.** Time-course of a perturbed trial. The trial follows the same baseline trial pattern described in 1C. However, the arm is perturbed perpendicularly to the straight line from the origin to the centre of the target, 1 to 3cm after the endpoint leaves the origin cue. Three separate factors are considered for perturbed trials: early/late, weak/strong, right/left.

The session consisted of 720 trials of two types: 2/3 baseline trials and 1/3 probe trials, which were pseudo-randomly interleaved. During baseline trials, subjects could perform their reaching movements without perturbation. During probe trials, the subject’s arm was mechanically perturbed in a direction perpendicular to the axis between the origin and the target sometime after the movement had been initiated and before the target had been reached. Each participant performed 10 blocks of 72 trials each in a single session (~1h15min). Each block consisted of 48 baseline trials (16 baseline repetitions of each distribution of reward) and 24 probe trials. There were 24 types of probe trials, reflecting each combination of the distribution of values (3-3, 5-1, or 1-5), the time at which the perturbation was applied (early or late after the movement onset), the direction of the perturbation (right or left) and the intensity of the perturbation (weak (3N) or strong (6N)). Each block contained one trial of each possible probed type (3×2×2×2=24). Trial order was counterbalanced and randomized both within and across blocks.

Real-time visual feedback of hand position was provided during the trial by a 1cm white dot on the screen, synchronized with the tip of the participant’s right-hand position on the experimental table. The time-course of each kind of trial is shown in FIG 1C-D. A baseline trial began when the origin was shown on the screen and the subject entered the cue. Approximately 1s later, the rectangular target and the distribution of reward were shown. After 500ms, the distribution of reward was removed. After a 1s interval, a GO signal was given when the origin cue vanished. The subject was instructed to perform a movement towards the position along the rectangle which they deemed most rewarding. If the subject left the origin before the GO signal was given, the experimental arrangement disappeared, and the subject had to wait until the regular trial duration of 7s elapsed before resuming the next trial. Correct target entry resulted in the rectangle turning green. After 500ms of holding position at the target, the target disappeared. This was followed by an inter-trial interval (ITI) of approximately 1s, the duration of which was dynamically calculated to obtain a fixed overall trial duration of 7s, to prevent participants from performing faster movements just to increase reward rate. The probe trials followed the same time-course of the baseline trials, with the exception of the mechanical perturbation, which was applied when the endpoint was either 1cm or 2.5cm away from the origin. At the beginning of each block, subjects were reminded that their goal was to maximize reward and that, during probe trials, they may have to change their mind to attain that goal.

### Muscle recordings

Electromyographic (EMG) activity was recorded from two flexors: pectoralis major (PEC), biceps long head (BIC); and two extensors: posterior triceps (TRI), posterior deltoid (DEL). EMGs were measured with disposable MT-130 surface electrodes (King Medical, ON, Canada), amplified (x5,000) and band-passed filtered (5-400Hz) by an 8-Channel Lynx-8 (Neuralynx, Bozeman, MT) and sampled at 1,000Hz by an acquisition card (National Instruments, Austin, TX) installed in a PC running Windows XP (Microsoft, Redmond, WA).

### Statistical tests

#### Analysis of Kinematics

Quantitative analyses of trajectories and velocities was performed with custom-written MATLAB scripts. First, we examined trajectories to determine the participant’s final choices, labeling them as Right or Left as a function of whether the movement endpoint lies on the right or left hemiplane defined by the line between the origin and crossing the long rectangle side perpendicularly (FIG 2A). We also labelled their initial choices as Initial Choice Right (ICR) or Initial Choice Left (ICL) as a function of whether the first 200ms of the path trajectory lie on the right or left side of that hemiplane. We first characterized baseline trials during which subjects could freely choose their trajectories in the absence of perturbation, to gain an insight on the subjects’ kinematics in the absence of perturbation. Next, to study how subjects changed their mind, we performed a comparative analysis during probe trials, examining the initial movement direction as well as the final movement endpoint.

**Figure 2.**
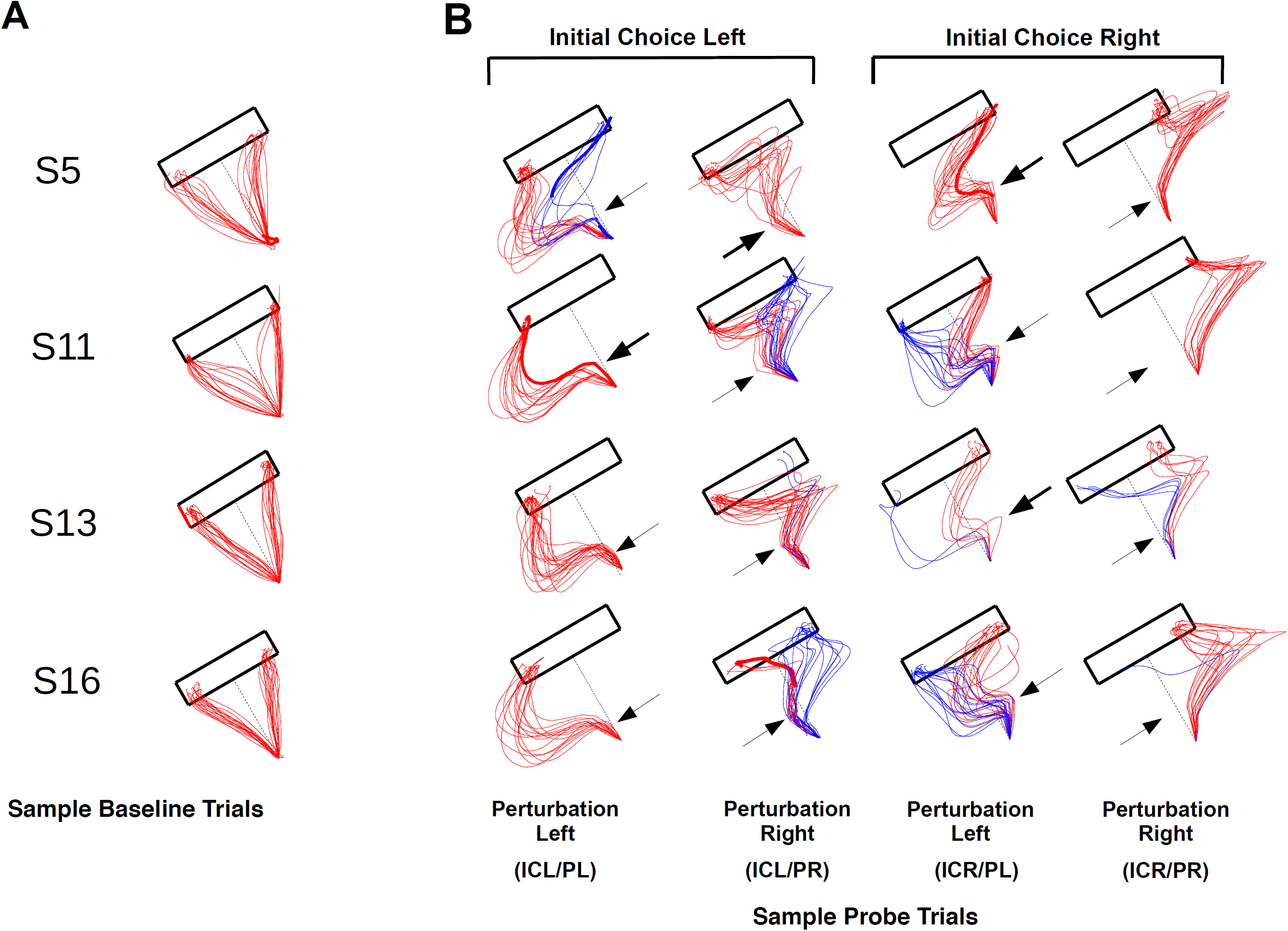
**A.** Typical baseline trajectories for subjects 5, 11, 13 & 16, during baseline trials. **B.** Perturbed trajectories for subjects 5, 11, 30 & 33, classified as a function of: 1) the initial choice direction (rightwards/leftwards) to the direction of motion; and 2) the direction of the perturbation (rightwards/leftwards). Blue trajectories indicate CoM trials, while red indicate non-CoM trials.

For our analyses, we used all subject data, and referenced their trajectories to the axis between the centre of the origin cue and the middle of the target rectangle’s long side. The origin of the reference system is the centre of the origin cue, with a positive y-axis in the direction towards the centre of the long rectangle side, and a positive x-axis from the origin towards the right side of the rectangle and parallel to its long side (FIG 1A). While the classification of baseline trials depended on their elected reaching target side alone (initial choice right/left), probe trials were classified, additionally, according to the time (Early/Late), intensity (Strong/Weak) and direction of the perturbation (Perturbation Left/Right), and, according to whether subjects changed their mind (FIG 2B). To assess the effect of the perturbations on the subjects’ trajectories, we first grouped trajectories into those that shifted to the target side opposite to their initial choice (CoM) trajectories, and those that remained on the same side (non-CoM). Second, we calculated the distance from each trajectory to the two lines defined between the origin and the bottom-right (DL1; Distance to L1) and bottom-left (DL2; Distance to L2) vertices of the rectangle long side (FIG 6).

Furthermore, we also calculated the radial and tangential velocities, as well as the tangential acceleration through differentiation for probe trials in each experimental case (FIG 3). Again, we assessed the effect of the change of mind by subtracting the differences between the velocity profiles during which there was a change of mind vs those in which there was no change of mind, for each type of probe trial.

**Figure 3.**
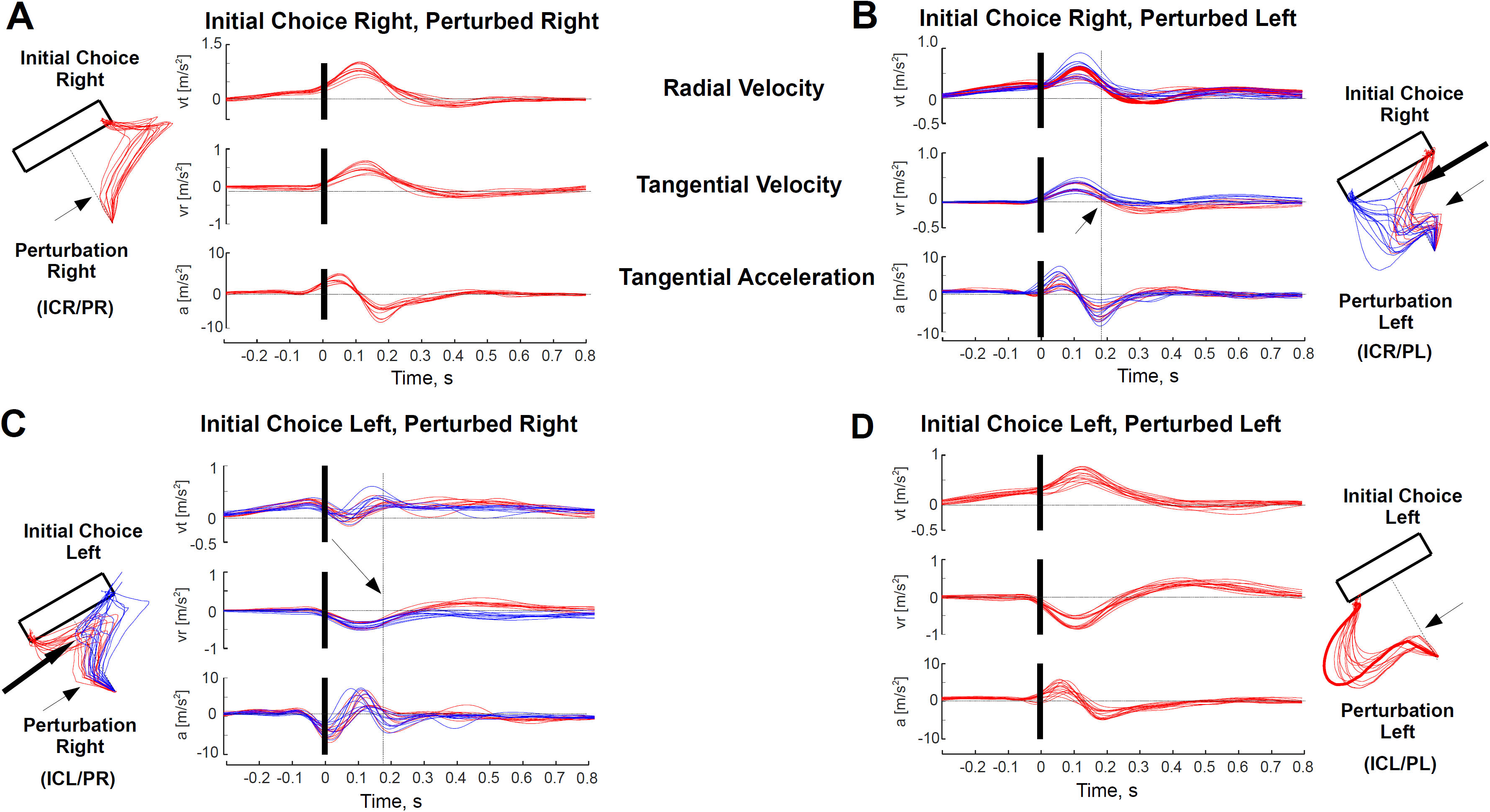
Radial velocities, tangential velocities, and tangential accelerations, aligned at the time of perturbation, for four kinds of trials, using data from S11: **A.** Initial choice right, perturbed right. **B.** Initial choice right, perturbed left. **C.** Initial choice left, perturbed right. **D.** Initial choice left, perturbed left.

#### Reward and Probability of Change of Mind

We first characterized the statistics of the changes of mind by estimating the p-parameter of a binomial random variable B(n,p) that would characterize the changes of mind, as a function of whether there was a CoM (PCoM =1) or not (PCoM=0). To that end, we counted the number of times each subject changed target side over the total number of probe trials. The resulting binomial p-parameter captures the probability of a CoM to occur.

We first estimated the binomial p parameter for each participant, and for each combination of the following three factors: initial choice right or left (ICR/ICL), perturbation direction (PR/PL), and the distribution of value (3-3, 1-5, 5-1). Second, we compared the p-parameter between cases in which the reward distribution was even (3-3) vs those cases in which opting for the target side opposite to their initial choice meant a large drop in predicted reward (5-1 or 1-5), as a function of whether the perturbation was towards the right (PR) or left (PL). Namely, we compared PCoM for the following four pairs of cases: ICR/PR, 3-3 vs 1-5; ICR/PL 3-3 vs 1-5; ICL/PR 3-3 vs 5-1; ICL/PL 3-3 vs 5-1. For each of the four cases, we assessed significance by means of a t-test on the difference of PCoM between cases of unequal reward (1-5 or 5-1) and equal reward (3-3).

Furthermore, to analyse the dependence of the PCoM on the remaining experimental factors, we pooled together trials regardless of DoV, and calculated the binomial p-parameter for each participant for each possible combination of perturbation time (T; early/late), perturbation intensity (I; weak/strong) and peak velocity prior to the perturbation (PV; slow/fast). T, I and PV were classified as Early/Late, Weak/Strong and Slow/Fast using median splits within the distribution of T, I and PV of each participant. We then used a mixed effects model and fitted a Generalized Linear Model (GLM) to the resulting p-binomial parameters against the three factors for each individual participant. Group significance was established via Bonferroni (and/or permutation tests) corrected F/t-tests on each of the β-regression coefficients. Significance was established if the probability of the null hypothesis was smaller than 5% (P<0.05).

#### The Temporal Unfolding of the Change of Mind

To study the temporal unfolding of the changes of mind, we performed an analysis of CoM vs. non-CoM trajectories during probe trials. Specifically, we focused our attention on differences on DL1 and DL2 trajectories between CoM and no-CoM trials, for each case of initial choice and perturbation direction: ICR/PR, CoM vs no-CoM; ICR/PL CoM vs no-CoM; ICL/PR CoM vs no-CoM; ICL/PL CoM vs no-CoM. For each participant, we aligned probe trial trajectories on the onset of movement and performed a sliding t-test on the distance metrics DL1 and DL2, between CoM vs. non-CoM trials, and calculated the two times along the path-trajectory at which CoM and non-CoM became significantly different with 95% and 99% probability.

We portrayed the temporal unfolding of the change of mind by means of two scatter plots of DL1 vs DL2, sampled at two times of interest along the trajectory: at the point of peak deviation post-perturbation, and at the time the difference between CoM and non-CoM trials in terms of DL1 (or DL2) reached significance at P<0.05. These scatter plots were also fitted with ellipses by means of Principal Component Analysis (PCA), aligning their axes with the dimensions of maximum variability, with radii equal to the square root of the corresponding eigenvalues. These scatter plots also served the purpose of characterizing the state of the motor apparatus at any given time.

#### The State of the Motor Apparatus

In addition of assessing the influence of reward on the PCoM, we also performed an analysis of the influence of reward on movement by assessing the differences in DL1 and DL2 as a function of reward at the peak deviation. Namely, we predicted that reward would influence movements either by magnifying responses (increasing vigour) (Guitart-Masip et al., 2011; Choi et al., 2014) or by increasing the subject’s concern for precision (diminishing arrival velocity) (Trommershäuser et al., 2003). For this analysis, we selected only non-CoM probed trials, and classified them according to choice and perturbation direction: ICR/PR, ICR/PL, ICL/PR, ICL/PL. Comparisons within each case were performed between predicted reward 3 vs 5 trials. The results are plotted separately for DL1 and DL2 as scatter plots and histograms as a function of reward case.

## RESULTS

### Choice Preferences

We first assessed the influence of reward on decision-making by calculating: 1) the target side preferences of each participant as a function of the reward distribution (3-3, 1-5, 5-1) during baseline trials; 2) the distributions of target arrival locations (Suppl. FIG 1A); and 3) initial directions with respect to the midline defined by the coordinate system in FIG 1A for each reward distribution (Suppl. FIG 1B). As expected, arrival distributions peaked around positions of the rectangle side that offered the largest amounts of reward for uneven distributions (1-5, 5-1), indicating that subjects’ choices were strongly biased by predicted reward. Furthermore, the initial directions showed that movements were most frequently directed toward those positions from the start.

### Analysis of Kinematics

FIG 2A shows some typical baseline trial trajectories during 3-3 reward distributions for four subjects. The choice of target side is equally as frequent towards the right or left side of the rectangle. FIG 2B shows typical trajectories for the same four subjects during probe trials. We classified probe-trial trajectories as a function of their initial direction, perturbation direction (right/left), and CoM/non-CoM occurrence (see METHODS). The task was designed to perturb with equal frequency towards the right and left side of the y-axis (FIG 1A) --- roughly perpendicular to the direction of movement, so that subjects could not anticipate the upcoming perturbation or its direction. We predicted that perturbations towards the side of the target opposite to the subjects’ initial choice were likely to yield a change of mind after the onset of movement. However, in addition to confirming this prediction (S11, S16 in FIG 2B), our results also show that a few changes of target side occurred when both the Initial Choice and the Perturbation directions matched (FIG 2B; S5, Initial Choice Left/Perturbation Left; S13 & S16, Initial Choice Right/Perturbation Right).

FIG 3 shows the tangential velocity and acceleration, and radial velocity traces for four types of probed trials, for a typical subject (S11). We aligned the traces at the time of perturbation. The four types of trials are: initial choice right, perturbed right (ICR/PR; FIG 3A), initial choice right, perturbed left (ICR/PL; FIG 3B), initial choice left, perturbed right (ICL/PR; FIG 3C), and initial choice left, perturbed left (ICL/PL; FIG 3D). A first visual analysis suggests that the first moment of divergence between non-CoM and CoM trajectories occurs approximately 170-80ms post-perturbation (FIG 3B & 3C). Incidentally, this matches the time of peak deceleration.

### Reward and Changes of Mind

CoMs may occur when the perturbation assists the movement towards the target side opposite to their initial choice (FIG 4A), and may be biased by the distribution of reward at that trial. One way of testing this is by calculating the PCoM for each reward distribution and to assess differences between cases. Specifically, we calculated the PCoM for each initial choice and perturbation direction case (ICR/PR 3-3 vs 1-5, ICR/PL 3-3 vs 1-5, ICL/PR 3-3 vs 5-1, ICL/PL 3-3 vs 5-1 --- FIG 4A), and obtained a mildly larger PCoM when the reward distribution is 3-3 over 1-5 and 5-1 (FIG 4B). These differences reach group significance when comparing PCoMs between inward (ICR/PL & ICL/PR) vs outward (ICR/PR & ICL/PL) perturbation conditions (FIG 4C; P=0.013). In other words, a CoM is most likely to occur during 3-3 trials and when perturbed inwards.

**Figure 4.**
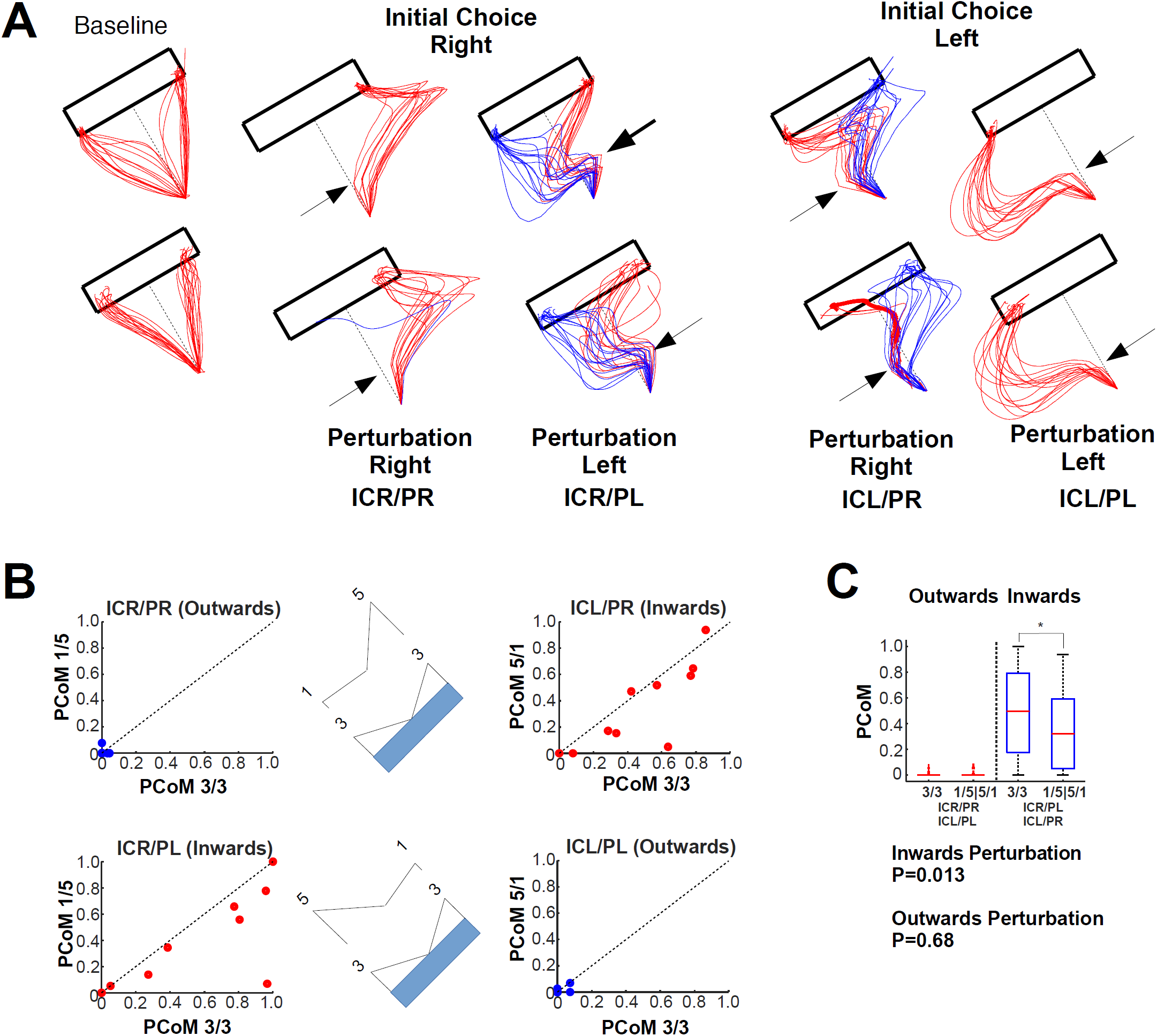
**A.** Baseline and probe trial trajectories (see FIG 3) for two subjects. **B.** Influence of Reward onto PCoM. Scatter plots of the average PCoM per subject when aiming at R=5 vs. R=3 in four cases: ICR/PR 3-3 vs 1-5; ICR/PL 3-3 vs. 1-5; ICL/PR 3-3 vs 5-1; ICL/PL 3-3 vs 5-1. **C.** Group PCoM average in the four cases described in B. Note that PCoM comparison in R=5 vs R=3 yields group significance in the case the perturbation is inwards (P=0.013).

In addition to reward, we also assessed the dependence of PCoM on non-explicit factors --- unknown to the subject at the time of the trial, such as the direction of the perturbation, its intensity and the hand velocity prior to the perturbation. To this end, we fitted a multivariate Generalized Linear Model to the binomial p-parameter values obtained per subject, and calculated for each combination of the aforementioned factors (FIG 5A-C). Group significance was obtained for perturbation intensity (F(1,12)=7.73; P=0.0066) and for the velocity (F(1,12)=22.21; P=8.60E-6), but not for the time of perturbation (F(1,12)=0.23; P=0.63) --- FIG 5C. In summary, changes of mind were more likely to occur after strong perturbations (FIG 5E) and during slow movements (FIG 5F).

**Figure 5.**
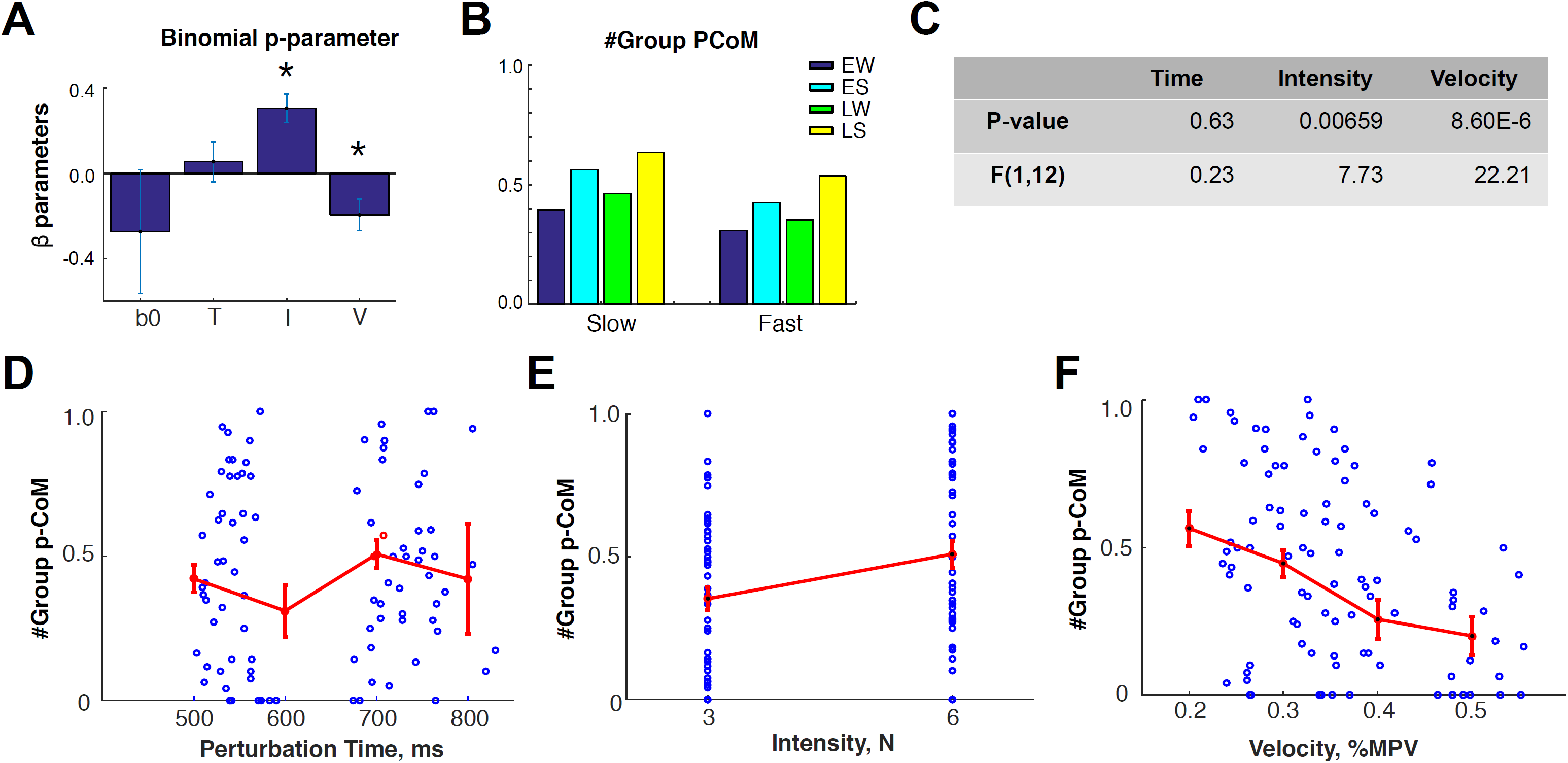
**A.** Group average regression coefficients a GLM on the p-binomial distribution parameter, calculated on an individual subject basis, as a function of three factors: Time of Perturbation, Intensity of Perturbation, Tangential Velocity. **B.** Group PCoM as a function of perturbation intensity and time of perturbation: Early-Weak (EW), Early-Strong (ES), Late-Weak (LW) and Late-Strong (LS). **C**. Group significance F and P-Values per factor (F(1,12)=7.728, p=0.00659) & Tangential Velocity (F(1,12)=22.21, p=8.60E-6). **D-E.** Group average effects for the Perturbation Time, Perturbation Intensity and Velocity at the time of Perturbation.

### State of the Motor System

We assessed the influence of the perturbation on the CoMs as the movement unfolded by measuring the distance between each probe trial trajectory, perpendicularly to the two lines used as reference axes for movements towards either rectangle side (from the origin to the right/left bottom vertices of the rectangle target, DL1 and DL2, respectively; see METHODS). Red and blue traces in FIG 6A & Suppl. FIGs 2-4, show the resulting path distances for four typical subjects (S4, S5, S11 and S32), alongside with the subject end-point trajectories, classified as a function of initially aimed target side (right/left), direction of perturbation (right/left) and CoM/non-CoM.

**Figure 6.**
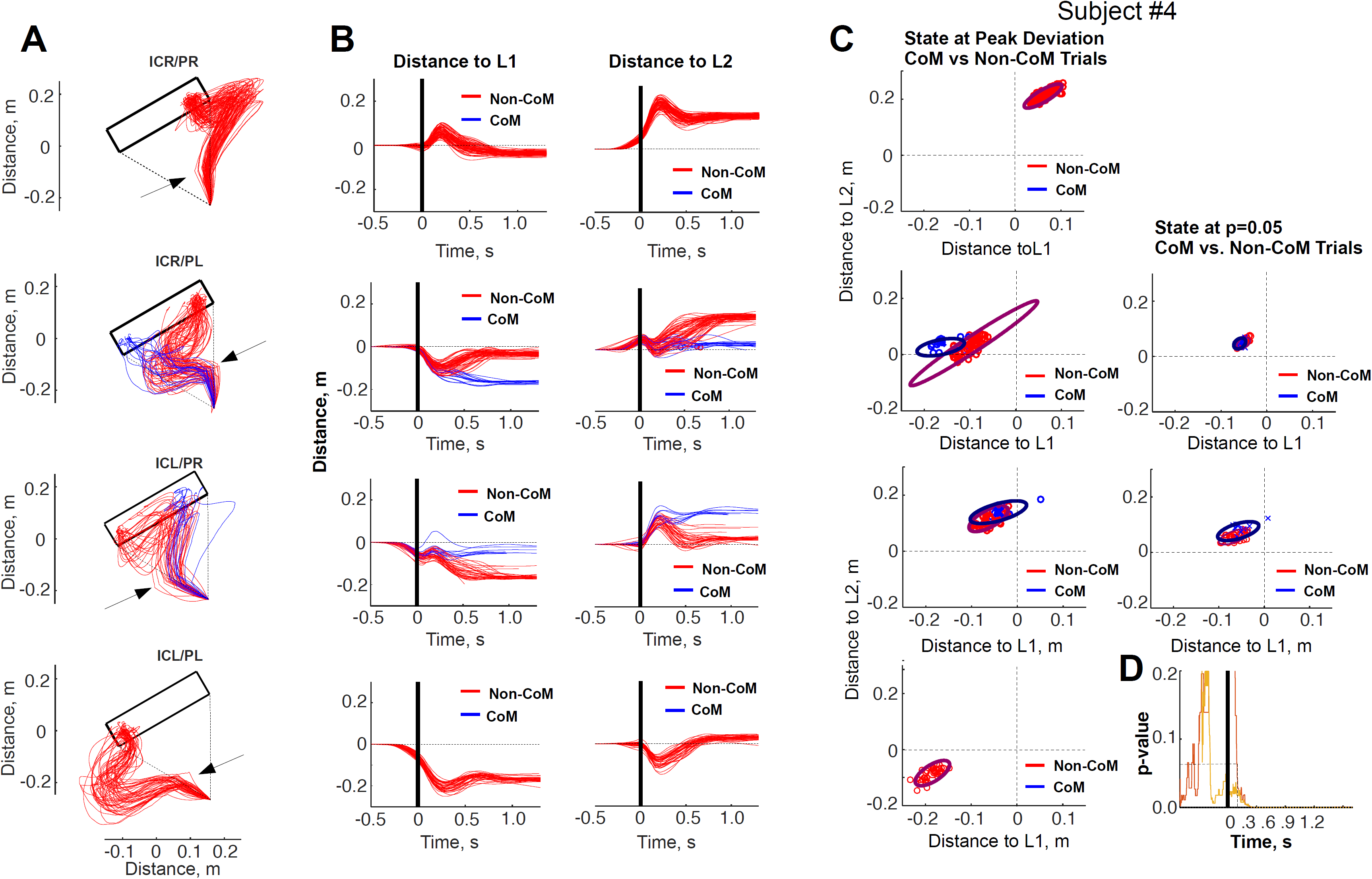
**A.** Probed trajectories for the cases of study: ICR/PR, ICR/PL, ICL/PR, ICL/PL for Subject#4. **B.** Distance to L1 and L2 in each of the four cases shown in A. Blue: CoM trajectories, Red: non-CoM Trajectories. **C.** State of the motor apparatus, characterized by DL1 and DL2 at the time of peak deviation, and at the time the difference between CoM and non-CoM trajectories became significant. **D**. P-value as a function of time, resulting from the sliding t-test (50ms window), performed between CoM vs non-CoM DL1 trajectories, during ICL/PR (red) and ICR/PL (gold).

Although the context in which the CoM may occur is generated by a mechanical perturbation that elicits a motor reflex in the opposite direction, the subsequent trajectory towards the opposite side of the target must be voluntarily enacted. Our hypothesis is that the gating of the change of mind depends on the state of the motor system at the time the perturbation is induced. If this were the case, we should observe significantly different states of the motor system in correspondence to perturbations that elicited CoMs versus those that did not.

To test this, we characterized the state of the motor system by the joint (bi-dimensional) path distance from each trajectory to L1 and L2 (DL1 and DL2), evaluated at the time of peak deviation. FIG 6C (and Supplemental FIGs 2C-4C) show this bi-dimensional state, plotted as x-y planar coordinates, for CoM (blue) and non-CoM (red) trials. We used an enveloping ellipse to capture data covariance. Since changes of mind were primarily elicited in trials where the direction of the perturbation and the target side initially aimed for were opposite, we restricted the CoM vs non-CoM comparison to those cases. We established statistically significant differences with a sliding t-test on DL1 and DL2 between CoM and non-CoM trials from movement onset until peak deviation (see METHODS). Remarkably, all subjects exhibited significant differences before the peak deviation (P<0.05).

### The Critical Timing for Changes of Mind

To establish the degree of anticipation with which the state of the motor system was different between CoM and non-CoM trials, we performed t-tests at 10ms intervals from movement onset until the time of peak deviation. With a P-value < 0.05 as our threshold of significance, we found that the times at which the state of the motor system was already significantly different ranged from 30 to 450ms after movement onset (avg = 153ms; Figure 6D & Suppl. Figure 5).

### The Effect of Reward on the State of the Motor System

We hypothesized that the state of the motor system, characterized by the relative distances DL1 and DL2, should vary as a function of predicted reward, making CoMs more likely to occur during 3-3 trials. To test this, we quantified the influence of reward on the state of the motor system, and assessed whether larger rewards yield faster movement initiations and longer deceleration phases towards the target. We compared our bi-dimensional path distance metric (DL1 & DL2) as a function of reward distribution during non-CoM probed trajectories, and restricted our analysis to trajectories aiming at target positions offering a reward of 3 (3-3) vs those aimed at reward 5 (5/1 or 1/5). Specifically, we performed the following four comparisons: ICR/PR 3-3 vs 1-5, ICR/PL 3-3 vs 1-5, ICL/PR 3-3 vs 5-1, ICL/PL 3-3 vs 5-1 (FIG 7). Their related trajectories are shown in FIG 7B.

**Figure 7.**
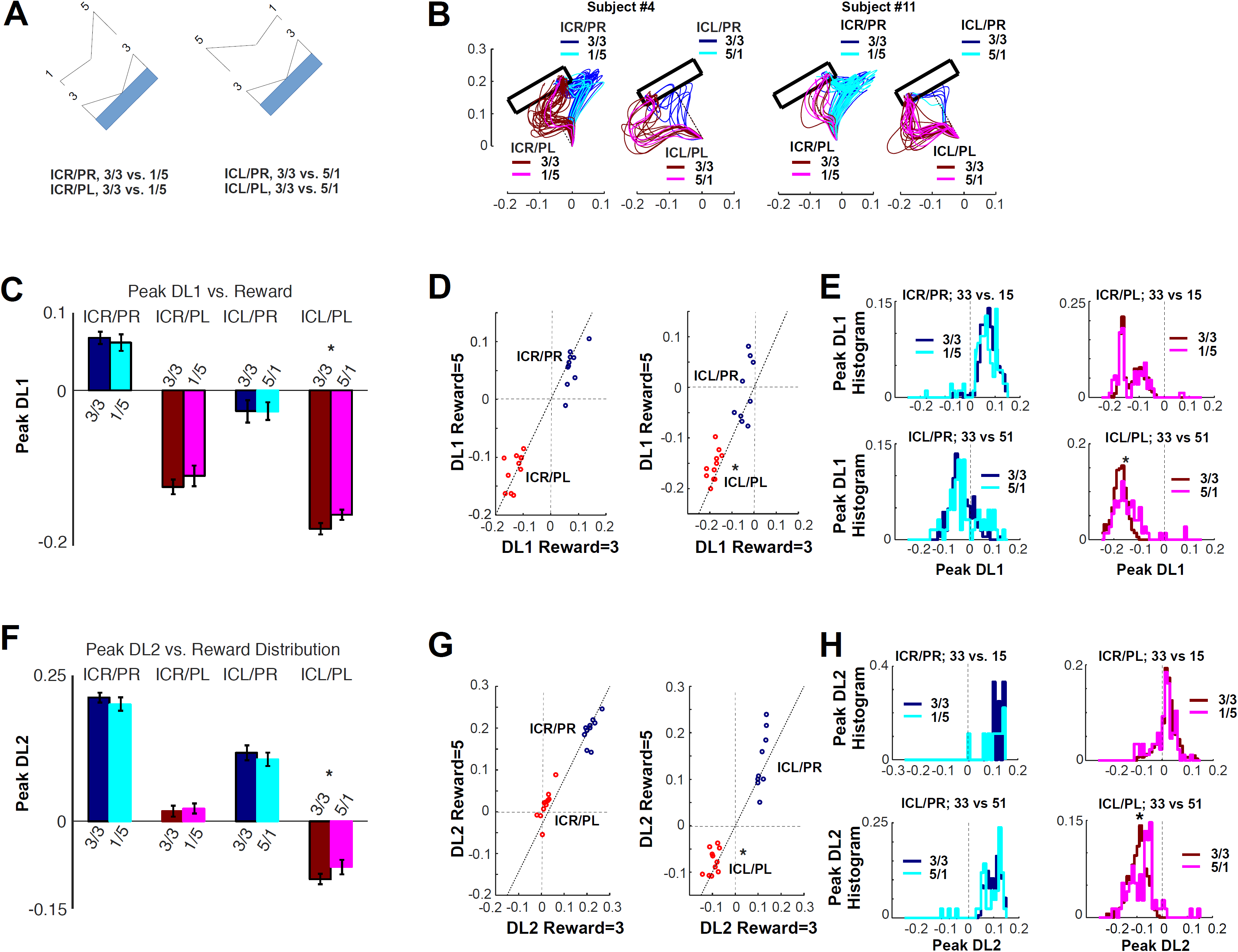
Effects of reward on DL1 and DL1 at Peak Deviation on probed non-CoM trials in four cases listed next: **A**. ICR/PR 3-3 vs 5-1, ICR/PL 3-3 vs 5-1, ICL/PR 3-3 vs 1-5, ICL/PL 3-3 vs 1-5. **B.** Sample trajectories for the cases described in A for subjects #4 and #11. **C.** Group average and standard error of the Peak DL1, for the cases described in A. **D**. Scatter plot of each subject’s average DL1, evaluated at R=5 vs R=3 trials. E. Group distributions of peak DL1, evaluated in the cases described in A. **F-H.** Same as C-E, but for DL2.

FIG 7C shows the group average and standard error of DL1 at the time of peak deviation, across probed trajectories for the four aforementioned reward comparisons. FIG 7D shows two scatter plots of the mean DL1 per participant, as a function of predicted reward --- 5 vs 3. If reward were not to exert any effect on the state, we should expect DL1 to assume equal values regardless of reward prospect, and to obtain average DL1 values aligned with the dashed diagonal in FIG 7D, and to obtain equal distributions of peak DL1 values for the four cases (FIG 7E). FIG 7F-H shows the equivalent comparisons for DL2.

Although the FIG 7C-E and 7F-H show trends consistent with that hypothesis, statistical tests report that the state of the motor system, was significantly different only in the case of initially aiming towards the left and being perturbed towards the left (ICL/PL). In this case, the distance (DL2) to the closest reference line – L2, exhibits a significantly smaller mean when aiming at 5 than when aiming at 3 (P=0.012). In other words, the effect of the perturbation is weaker when aiming at 5 than when aiming at 3, strongly suggesting that in the certainty of an absent more desirable option, the commitment is stronger. While a similar trend is also visible in the ICR/PL and ICL/PR cases, the size of the effect does not reach group significance (P=0.072; P=0.14).

## DISCUSSION

Traditional models describe decision-making as a sequential process of deliberation followed by commitment, all of which precede the planning and execution of the chosen action. However, in ecologically valid conditions, it is often useful to initiate action before the deliberation phase is complete, and to revise the initial plan along the way, as a consequence of (for example) changing prospects or motor costs (Cisek and Pastor-Bernier, 2014b; Lepora and Pezzulo, 2015; Domínguez-Zamora and Marigold, 2019). This *embodied choice* perspective suggests that the deliberation process has to remain flexible even after action initiation, to permit re-evaluating the initial choices – and “changing mind” when necessary. While changes of mind during action execution have been observed empirically (Resulaj et al., 2009; Song and Nakayama, 2009; Barca et al., 2012), it is not clear whether they reflect a truly deliberative process, and whether they are sensitive to economic values (i.e., the values of the selected and unselected offers) and situated aspects of the choice (e.g., the state of the motor system during the choice, and the costs of revising the initial plan).

Here we studied these economic and motor-related determinants of changes of mind during decision-making, by exploiting the fact that changes of mind can be triggered externally by perturbing the motor apparatus during the choice (Nashed et al., 2014). We designed a task in which participants had to select a reaching path trajectory from an origin to a wide rectangular target, where the reward was distributed non-uniformly as a function of the arrival endpoint. Rewards could be even (3-3) or uneven (1-5/5-1) at the two target sides. Critically, we applied mechanical perturbations after the choice was made, at different phases during the movement and in different directions, sometimes toward the lower-valued side of the target.

Our results show that, as expected, participants facing a choice between two target sides offering different rewards initially selected the direction of movement offering the highest prospect; whereas participants facing a choice between two regions offering the same prospect made their selection with approximately equal frequency. After a perturbation was applied, participants altered their initially selected target side most frequently when the change did not result in a reduction of reward, the perturbation was more intense, the movement was slower and the distance from the initially selected target was greater. These results indicate that changes of mind were influenced both by the predicted reward associated with the action and the state of the motor system at the time of the perturbation.

These findings have two main implications. First, they provide supporting evidence for the notion that deliberation continues and remains flexible after movement onset, possibly because unselected potential actions are not completely suppressed or discarded (Lepora & Pezzulo, 2015). From a neurobiological perspective, the *affordance competition hypothesis* suggests that situated choices are resolved through a biased competition between neuronal populations corresponding to potential actions that implement the competing choices, as observed in the monkey premotor cortex (Cisek, 2007; Cisek and Kalaska, 2010; Pezzulo and Cisek, 2016). Our findings suggest that when there is a chance to revise the initial decision, the competition between potenital actions is not completely settled before action initiation, possibly because the initially unselected neuronal population retains some sub-threshold activation, and can take control afterwards.

Second, our findings suggest that decision-makers never completely commit to their initial choice during situated decisions. Rather, they have a variable degree of commitment to their initial choice, which depends on the relative value of the offers and on the state of the motor system. The pattern of results we observed suggests that the commitment to the initial choice is weaker when the decision is between even-valued choices, which makes changes of mind more likely. Furthermore, the commitment is stronger if the initially selected action leads to higher rewards, which makes the initial choice more resistant to external perturbations. Furthermore, our results also suggest that for a change of mind to occur, the state of the motor system has to be within a specific range, and that the external perturbation can act as a trigger for a change of mind only when departing from those states. Note that one can interpret the state of the motor system both as a *reflection* of a centrally computed degree of commitment (e.g., because one is committed, one decides to move faster) or as a *cause* of commitment (e.g., the action can be fast purely due to variability, but if it is fast, it is less prone to changes of mind). These two hypotheses remain to be disentangled in future studies.

More generally, our findings fit within a growing body of work suggesting that during natural behavior, decision-making and movement planning unfold together, in an integrated fashion, within highly distributed circuits spanning what have traditionally been considered purely cognitive versus sensorimotor regions of the brain (Shadlen et al., 2008; Cisek and Kalaska, 2010). Of course, in some conditions, such as abstract decisions between stable options without the pressure to act (e.g. choosing a chess move, deciding about a house to buy), the processes of outcome valuation and action control will occur at very different moments in time, each engaging only a restricted subset of the relevant neural mechanisms. Nevertheless, the organization of these mechanisms evolved for a different type of situation, regularly encountered during natural behavior, in which animals must make decisions even during ongoing sensorimotor activity.

## Acknowledgements

We wish to thank Ricard Beyloc-Jerez for producing the experimental setup figure. This work was supported by Marie Skłodowska-Curie Grant Scheme IF-65652 to IC, the Spanish Project PID2019-105093GB-I00 (MINECO/FEDER, UE) and CERCA Programme of the Catalan Government to IC, the European Union’s Horizon 2020 Framework Programme for Research and Innovation under the Specific Grant Agreements No. 785907 and No. 945539 (Human Brain Project SGA2 and SGA3) to IC and GP, a grant to GP from the European Research Council (Grant Agreement No. 820213, ThinkAhead), and a Natural Sciences and Engineering Research Council grant RGPIN/05245 to PC.

## Supplemental Figures

**Supplemental Figure 1 (related to Fig 2). A.** Distribution of endpoint positions at arrival to the target as a function of reward distribution: 3-3 (red), 5-1 (green), 1-5 (blue) for each individual subject, during non-perturbed (baseline) trials. **B**. Distribution of arrival positions as a function of value distribution: 3-3 (red), 5-1 (green), 1-5 (blue) for each individual subject, during non-perturbed (baseline) trials.

**Supplemental Figure 2 (related to Fig 6).** Same as FIG 6, for S5.

**Supplemental Figure 3 (related to Fig 6).** Same as FIG 6, for S11.

**Supplemental Figure 4 (related to Fig 6).** Same as FIG 6, for S32.

**Supplemental Figure 5 (related to Fig 6D).** P-Value of the t-test calculated between trajectories in which each subject changed his/her mind and the trajectories in which the subject stuck to his/her original choice after a perturbation. The comparison was performed by first calculating the distance between each trajectory and the straight path defined between origin and the right- or left-bottom vertex of the target. The t-test is calculated at each time step between both distribution of distances, and is calculated for each subject individually. **A**. P-Values as a function of time for each individual subject. **B**. Time in ms at which the P-Value for each subject became significant at 0.05 (black bars) and 0.01 (white bars).

