## Supplementary figures and images for "Changes of mind after movement onset: a motor-state dependent decision-making process"

### Supplementary Material

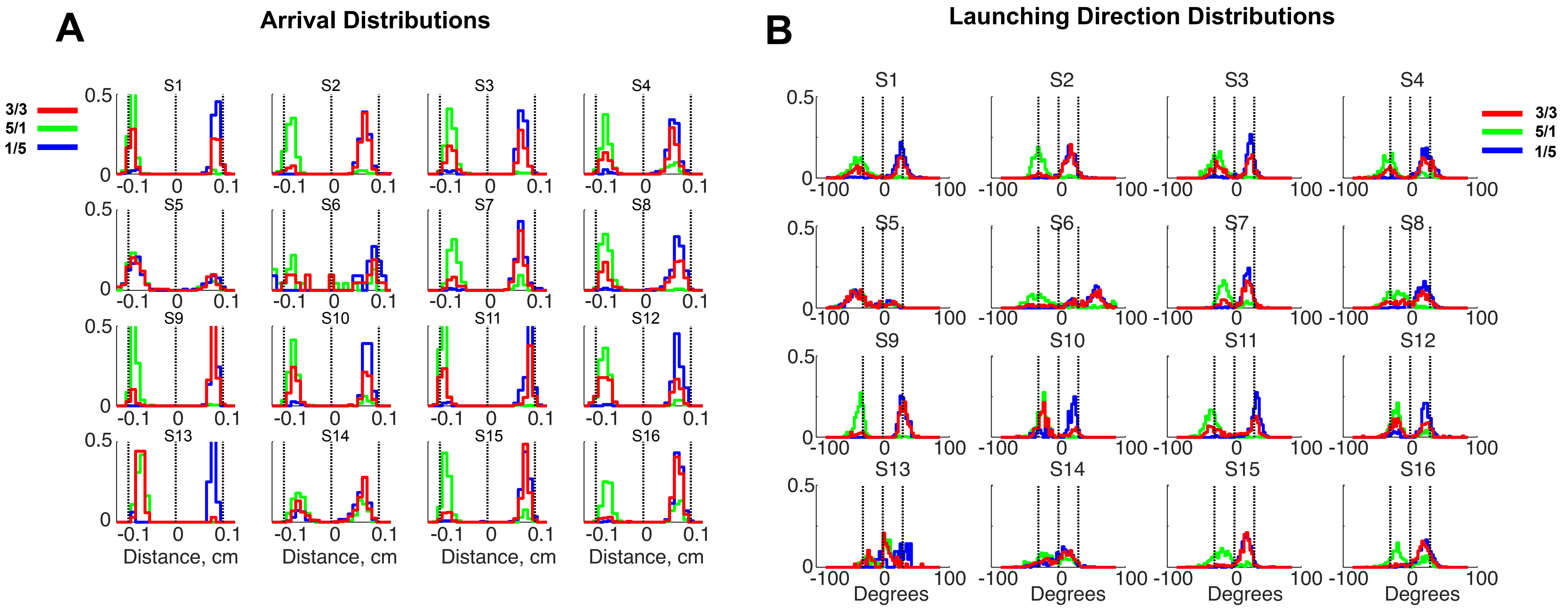

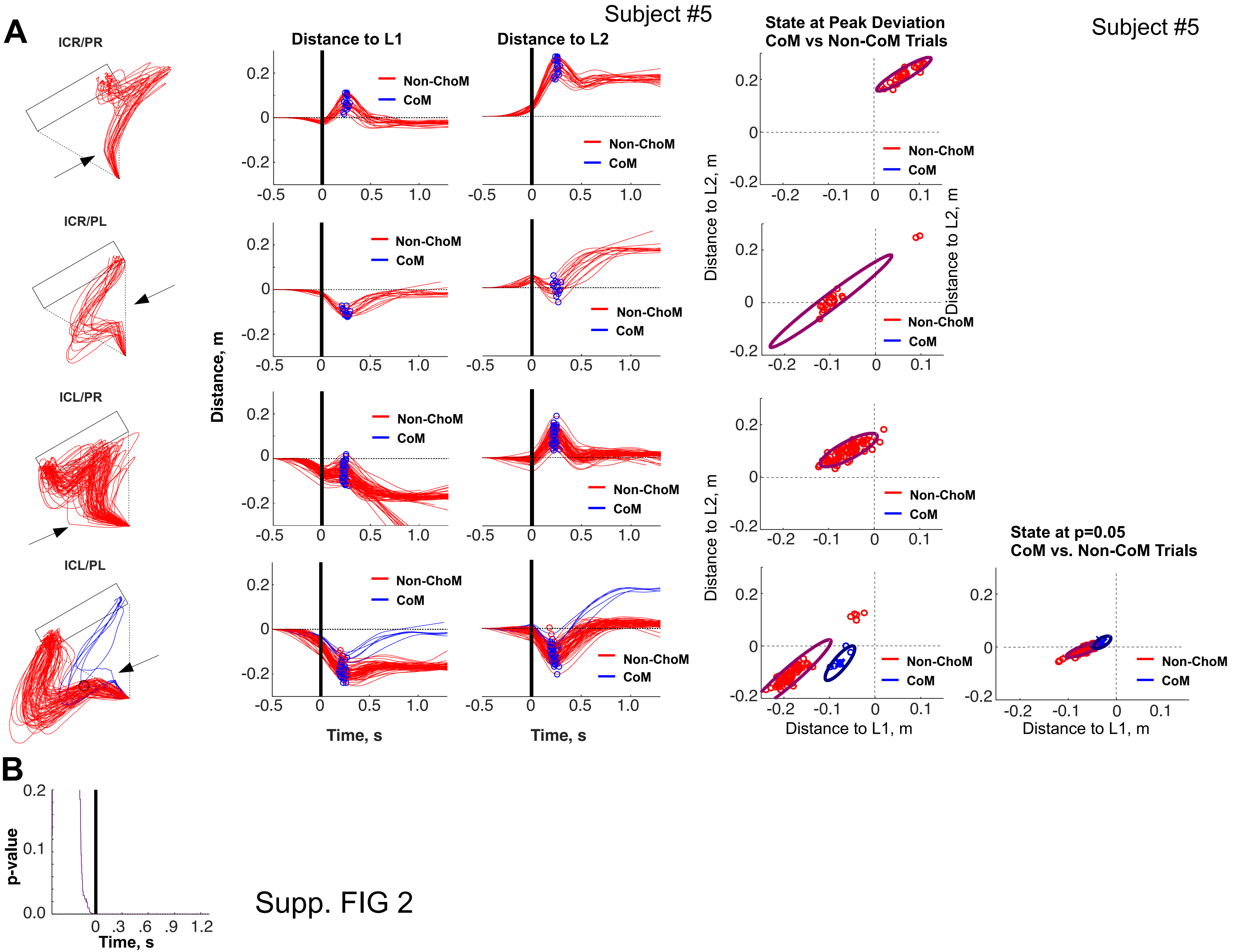

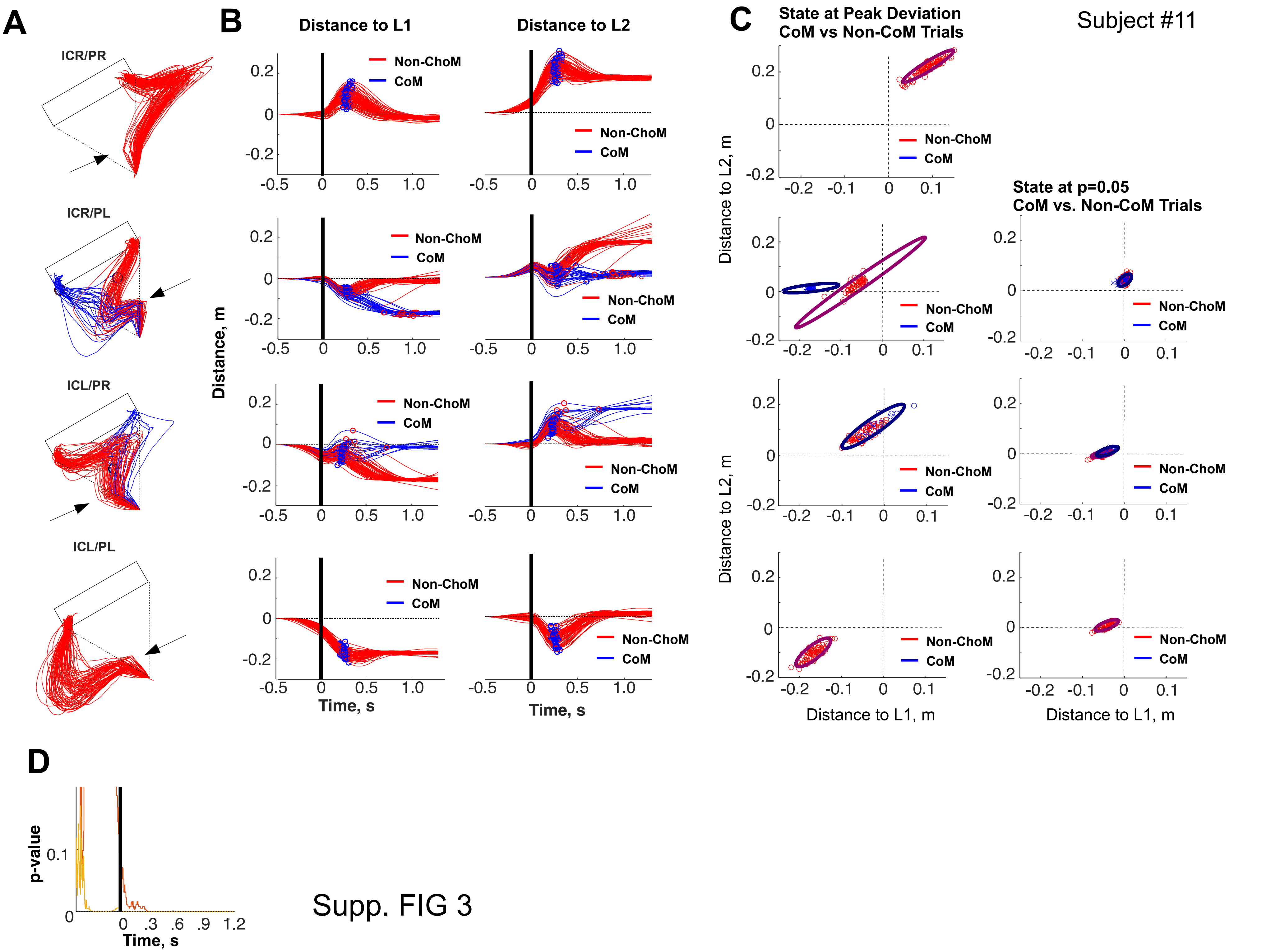

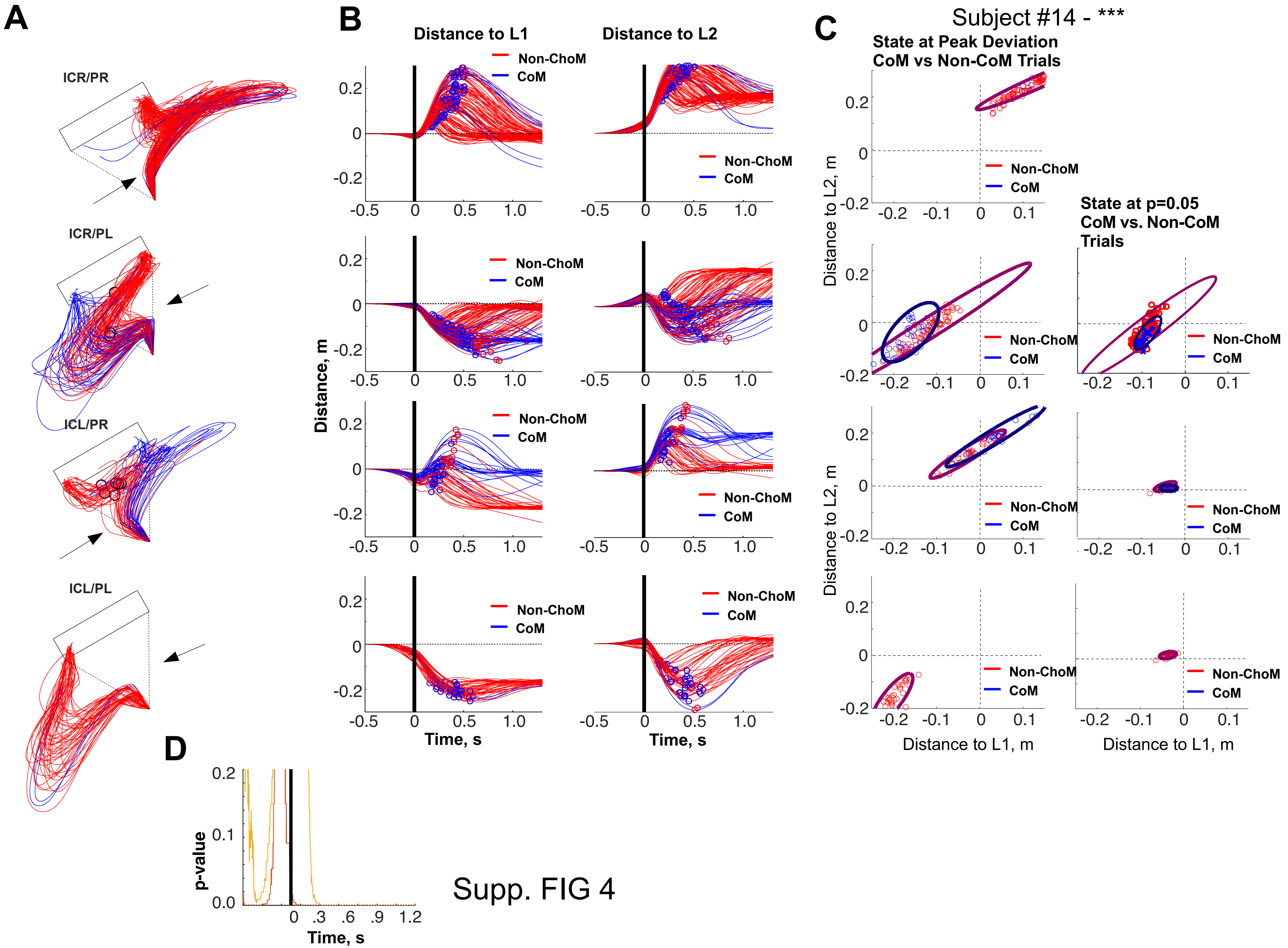

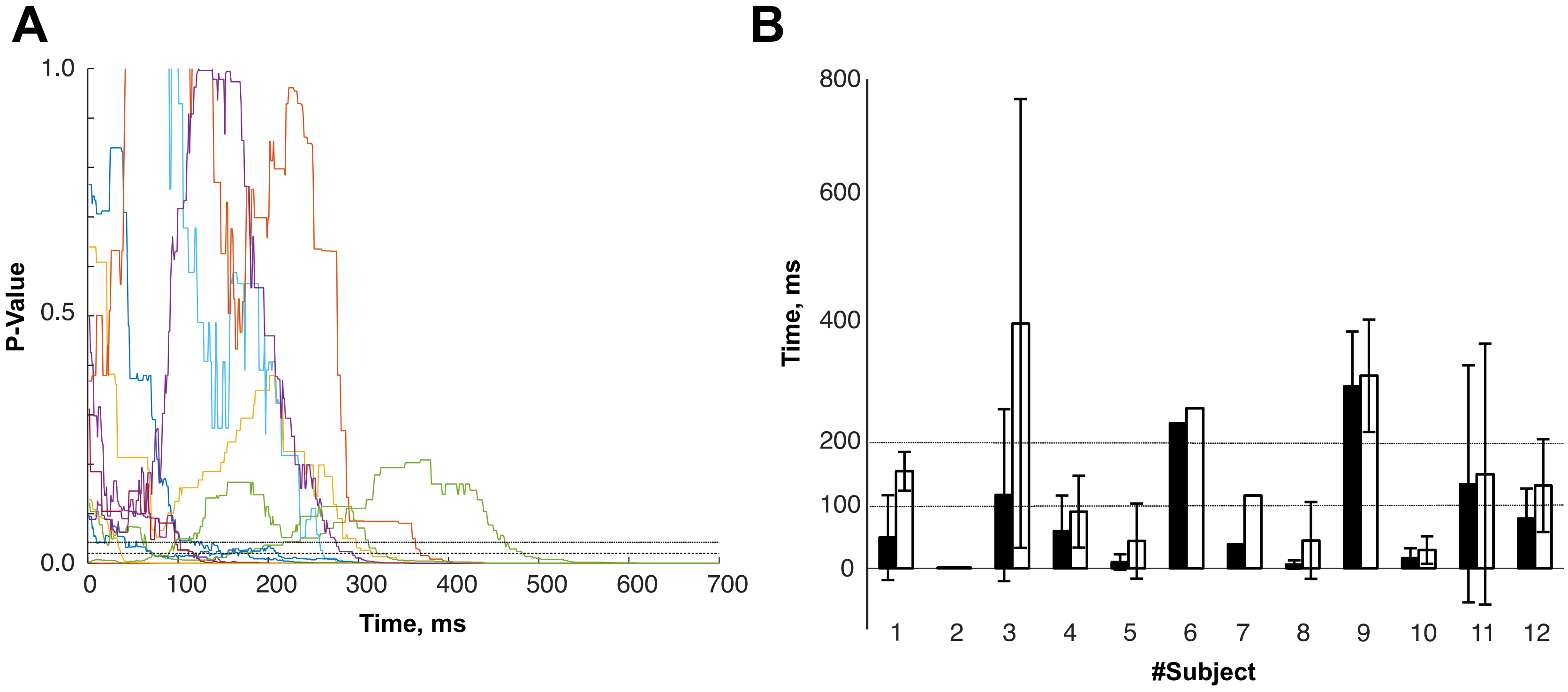
